# Canalization masks high levels of cryptic epigenetic and transcriptional plasticity in butterflies

**DOI:** 10.64898/2026.09.29.754141

**Authors:** Noah K. Brady, Karin R. L. van der Burg, Robert D. Reed

**Author notes:** Corresponding author: Noah K. Brady. These authors should be considered co-first authors (Noah K. Brady, Karin R. L. van der Burg). Department of Pathology, University of Southern California Keck School of Medicine, Los Angeles CA, 90033, U.S.A.

## Abstract

Phenotypic plasticity is a common adaptation to heterogeneous environments. While it is well documented that trait plasticity can rapidly evolve, little is known about the developmental mechanisms that underlie variation in plasticity. Here, we compare diverged populations of the common buckeye butterfly (*Junonia coenia*): one population with canalized wing patterns that are insensitive to environmental conditions and another population that has distinctive seasonal color patterns. In common garden trials we were surprised to find dramatically elevated levels of transcriptional and chromatin accessibility plasticity during wing development of the canalized population compared to the seasonally plastic population. Further, using machine learning analysis of accessible chromatin sequences, we observed a broad heterochronic shift in motifs enriched in plastic putative regulatory elements to earlier stages of development in the canalized population, which we speculate may cause phenotype-related gene expression to move outside a wing color critical period. Together, our data lead us to propose a model where cryptic genetic processes are masked by a shift in chromatin remodeling, which in turn could produce a capacitor-like effect to maintain developmental genetic variation in the canalized population.

## Introduction

Phenotypic plasticity - when an organism’s phenotype changes in response to environmental cues - is a widespread phenomenon in nature. Plasticity increases phenotypic variance, affects population persistence, and consequently can shape adaptation (Charmantier et al. 2008; Scoville and Pfrender 2010; Ghalambor et al. 2015; Fox et al. 2019; Campbell-Staton et al. 2021). Despite an appreciation for the role of plasticity in evolution, it is challenging to integrate concepts into a unified model without understanding the mechanisms that imbue traits with environmental sensitivity and how such mechanisms evolve to shape adaptive trait variation (Pigliucci 2001; West-Eberhard 2003; Beldade et al. 2011; Sommer 2020).

To identify developmental mechanisms that facilitate trait plasticity, gene expression is often measured across environmental conditions. In many instances, there is a positive relationship between the number of genes that are differentially expressed between environmental treatments and the degree of physiological or morphological plasticity (e.g. killifish, sticklebacks, honeybees, locusts, and beetles) (Alaux et al. 2009; Morris et al. 2014; Shaw et al. 2014; Casasa et al. 2020; Foquet et al. 2021) However, in some species that are adapted to highly variable environments, less environmentally sensitive gene expression is associated with more physiological plasticity, particularly in components of the adaptive stress response pathway (e.g. flies, corals) (von Heckel et al. 2016; Bay and Palumbi 2017). In these cases, there is still some plastic gene expression, but it is a component of a homeostasis mechanism that buffers environmental inputs. Thus, it remains unclear whether there is a general expectation about what the association between gene expression plasticity and trait plasticity should be.

Beyond gene expression, even less is known about the deeper epigenetic correlates of plasticity. Phenotypic plasticity relies on epigenetic processes – such as changes to chromatin accessibility - that regulate gene expression to produce alternate phenotypic outputs from a single genotype (Lyko et al. 2010; Ozawa et al. 2016; Vogt 2022; Li et al. 2025). These epigenetic changes can occur at cis-regulatory elements (CREs), the nodes of gene regulatory networks that control which transcription factors activate or repress gene expression (Buecker and Wysocka 2012). CREs must be unwound from nucleosomes by chromatin remodeling enzymes or pioneer factors in order for their protein-binding motifs to be bound by transcription factors (Lee et al. 2004; Klemm et al. 2019). Changes to CRE accessibility drive temporal and spatial patterns of development and can be epigenetically modified by environmental signals (Niederhuber and McKay 2021; Lowe et al. 2022; Hu et al. 2024; Tian et al. 2025). Therefore, epigenetic control of CRE accessibility could play a critical role in the development of plastic morphological traits, yet these processes remain poorly understood.

Adaptive developmental plasticity, when environmental cues during development produce traits that increase fitness in one condition versus another, is widespread throughout the insects (Lüscher 1960; Emlen 1994; Nespolo et al. 2009; Simpson et al. 2011; Song et al. 2017; Abouheif 2021). In particular, many butterflies and moths exhibit wing pattern plasticity as an adaptation to seasonal environments (Kingsolver and Wiernasz 1991; Otaki et al. 2010; Halali et al. 2021). Species within the nymphalid tribe Junoniini illustrate this phenomenon by changing multiple aspects of wing morphology between seasons, including pigmented colors, structural colors, color patterns, wing size, and wing shape (Clarke 2017). The North American common buckeye, *Junonia coeni*a, typifies this genus, varying from a light tan summer morph to a dark red autumn morph (Fig. 1A) (Smith 1993). This change is cued by seasonal variation in temperature and photoperiod, with autumn conditions delaying the timing of an early pupal pulse of the molting hormone ecdysone, a shift that is necessary for the development of the red morph (Rountree and Nijhout 1995). Experimental work demonstrated that the red autumn morph could rapidly become fixed in a population via selection on multiple loci, including a recently discovered long non-coding RNA (van der Burg. et al. 2020; Fandino et al. 2024). But the molecular changes that generate variation in the degree of plasticity between wild populations of *J. coenia* remain unknown. Research on this species has focused on a North Carolina population, which exhibit the aforementioned morphological plasticity. However, Southern California populations display almost no variation in color phenotype associated with the seasons, producing a fixed tan, summer-like phenotype regardless of environmental condition (Fig. 1B,C) (Daniels et al. 2012).

**Figure 1.**
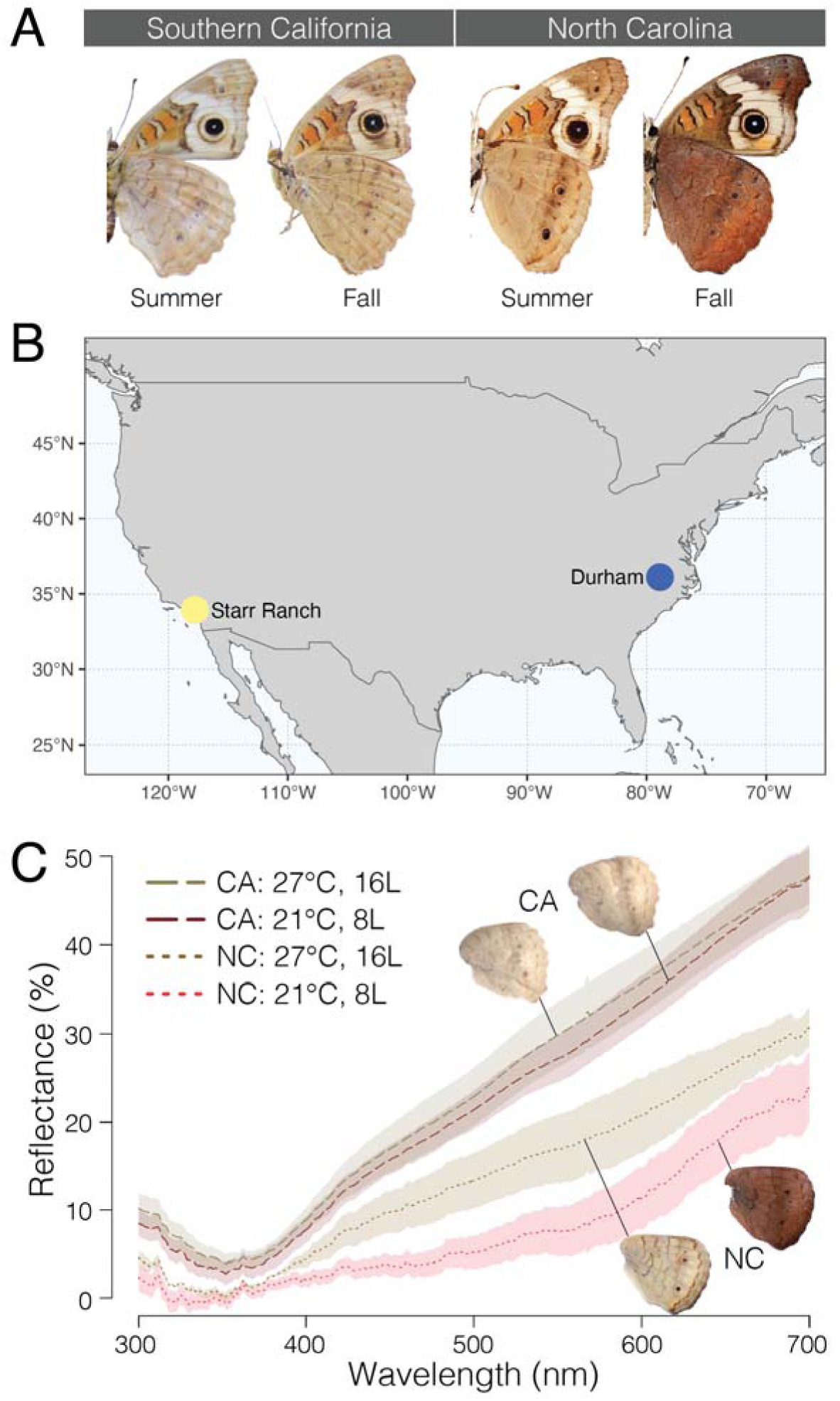
J. Coenia have diverged phenotypic plasticity of wing patterns between North Carolina and California. *A*) the dorsal color of *J. coenia* wings changes between seasons in North Carolina but not California B) Map of *J. coenia* study sites C) California *J.* coenia wings have higher reflectance scores (corresponding to being more tan) than North Carolina, where summer morphs have higher scores than fall morphs **Alt text**: comparison of wing morphology between study sites. A is a series of photos of J. coenia dorsal wings, showing color differences between seasons and populations. B is a map showing where animals for this study were collected. C is a plot of wing reflectance for summer and fall morphos of each population, with % reflectance on the Y axis and wavelength on the X axis.

Here, we leverage this variation, along with common garden experiments, measures of gene expression, and quantification of chromatin accessibility, to assess how canalization is associated with changes to developmental mechanisms underlying trait plasticity. We characterized variation in the number of plastically expressed genes across wing development in both populations and found a significantly elevated amount of transcriptional plasticity in the canalized population as compared to the morphologically plastic population. We also quantified the extent of environmentally induced changes in chromatin accessibility across development in both populations and again found elevated levels of plasticity in the canalized population. Many of these epigenetic loci neighbor genes that have diverged plastic expression, suggesting that these regions could include CREs that modulate gene expression plasticity. Furthermore, we identified a heterochronic shift in a distinct set of transcription factor motifs that are enriched in plastic putative regulatory elements. Our findings lead us to propose a model in which the plasticity of traits can evolve by shifting critical environment-transducing mechanisms in or out of developmental windows in which phenotypes are specified.

## Methods and Materials

### Animal collection and rearing

We collected wild *J. coenia* from two populations: Starr Ranch, in Trabuco Canyon, Southern California and a county park outside of Durham, North Carolina. The Southern California population we sampled is also described by some authors as *Junonia coenia grisea* or *Junonia grisea*(Gemmell and Marcus 2015). We reared wild-caught animals at either 27, 16 h light (summer-like) or 19 and 8 h light (fall-like), using an artificial diet supplemented with dried *Plantago lanceolata* (Nijhout, H. F. 1980). We used tissues from the first generation of these animals for RNA-seq and ATAC-seq experiments.

### Reflectance Measurements

We used Ocean Optics SpectraSuite software to take measurements of mid-ventral hindwings (between M3 and Cu2 veins) from 3 females and 3 males using an Ocean Optics USB2000 spectrophotometer equipped with a R200-7-UV/VIS reflection probe and a strobe light as a light source.

### RNA extraction and sequencing

We collected hindwing tissue from three replicates at four stages of development: late fifth-instar, pre-pupation, 72 h post-pupation, and 144 h post-pupation. We determined larval and pre- pupal stages using vein markers, pigmentation, and imaginal disc shape (Reed et al. 2007). At each stage, we dissected tissues in cold PBS, immediately transferred them to Trizol, and extracted RNA with the Invitrogen RNA PLUS kit. We prepared RNA libraries using the NEBNext Ultra RNA Library Prep Kit. We sent these libraries to the Genomics Facility at the Cornell Biotechnology Resource Center (BRC) for sequencing, which was carried out on an Illumina NextSeq500 on 2 x 36 bp lanes to a minimum depth of 20 million reads.

### Differential expression analysis

We filtered raw reads to remove low quality, multi-mapped, and duplicate reads and aligned pre- processed reads to the *J. coenia* (V2) transcriptome using Salmon (Patro et al. 2017; van der Burg et al. 2019). Following alignment, we carried out all further analyses using the R package DESeq2 (Love et al. 2014). We performed two primary analyses to quantify differences in log fold change of all expressed genes at each stage of development that we sampled: a) between environmental treatments within each population; and b) between populations but within each environmental treatment This program normalizes transcript counts based on library size among samples and fits a generalized linear model with a negative binomial distribution for each gene. After GLM fit, DESeq2 uses a Wald test to compare whether log2 fold changes between contrasts are significantly different from zero. For this test, we considered FDR adjusted p-values <0.05 to be differentially expressed.

### ATAC sequencing

For our ATAC-seq experiments, we collected hindwing tissues from three individuals (three replicates) at the same four stages of development as our RNA-seq experiment: late fifth instar, pre-pupal, 72 h after pupation, and 144 h after pupation (North Carolina fall-like samples from 5^th^ instar and 144 h had two replicates). We extracted nuclei, performed TN5 transposition, and prepared libraries following a modified version of the Buenrostro et al (Buenrostro et al. 2015) protocol that is tailored to butterfly wing tissues (Lewis and Reed 2019). We then sequenced these libraries to a depth of 20 million reads on 2x36 bp lanes of an Illumina NextSeq500 at the Genomics Facility of the Cornell Biotechnology Resource Center (BRC).

### ATAC-seq analysis

We aligned ATAC-seq reads to the *J. coenia* genome (V2) with bowtie2 after removing duplicate and multi-mapping reads. To assess sequencing quality and depth, we performed QC on fastq and bam files with FastQC. We called peaks from the aligned reads with MACS2, and assessed the signal to noise ratio of these calls using the fraction of reads in peaks (FRiP) using the program featureCounts (Zhang et al. 2008; Liao et al. 2014). We performed all downstream differential accessibility analyses in R. We assessed library complexity and fragment size distributions with the program ATACSeqQC (Ou et al. 2018). Finally, we quantified differentially accessible peaks using the R package DiffBind (Stark and Brown 2011).This package counts the number of aligned reads that are present within each peak that is shared among all samples and then uses DESeq2 to calculate log2 fold changes between groups. We used DiffBind to make pairwise comparisons of the number of peaks that changed accessibility between environmental treatments and between populations at each stage.

We used ChromBPNet to predict *de novo* motifs in open chromatin regions at each stage of development (Pampari et al. 2025). ChromBPNet is a deep learning program that corrects for the positional bias introduced by the TN5 enzyme and then predicts accessibility at a base-pair level. We used the count and profile scores from this program as input into TF-Modisco to aggregate the base-pair-level scores into motifs (Shrikumar et al. 2018). We merged motifs with high similarity and collapsed similar motifs. We note, however, that there were multiple butterfly motifs that matched with the same *Drosophila* motif; we treat them as distinct motifs and suggest that they could represent either different, context-specific binding sites of the same protein or binding sites for closely related proteins that lack known *Drosophila* homologs. Finally, we used the MEME suite of tools in R to run Fischer exact tests of enrichment to identify motifs that are disproportionately present in plastic ATAC peaks compared to all ATAC peaks (Bailey et al. 2015; Nystrom and McKay 2021).

## Results

### Transcriptional plasticity is greater in a morphologically canalized population

The degree of wing color plasticity differs between North Carolina and Californian populations of *J. coenia*. We sought to test whether the degree of morphological plasticity is correlated with the degree of gene expression plasticity by rearing F1 generations of wild-caught animals from both populations under warm, summer-like conditions and cool, fall-like conditions and then quantifying transcript abundance with bulk RNA-sequencing of developing wing tissue. All samples passed QC filters, with 65% of reads mapping to the *J. coenia* transcriptome (Table S1) and samples generally clustering by stage and condition (Fig. S1). Both populations exhibited significant levels of environmentally responsive gene expression (Fig. 2A, S3, S4), and this gene expression plasticity was highly variable across developmental timepoints (between 76-5114 differentially expressed genes, depending on stage; Table S3, S4). Surprisingly, there were many more genes differentially expressed between rearing treatments in the morphologically canalized California population versus the plastic North Carolina population (Fig. 2B; California: 6503; North Carolina: 2449).

**Figure 2.**
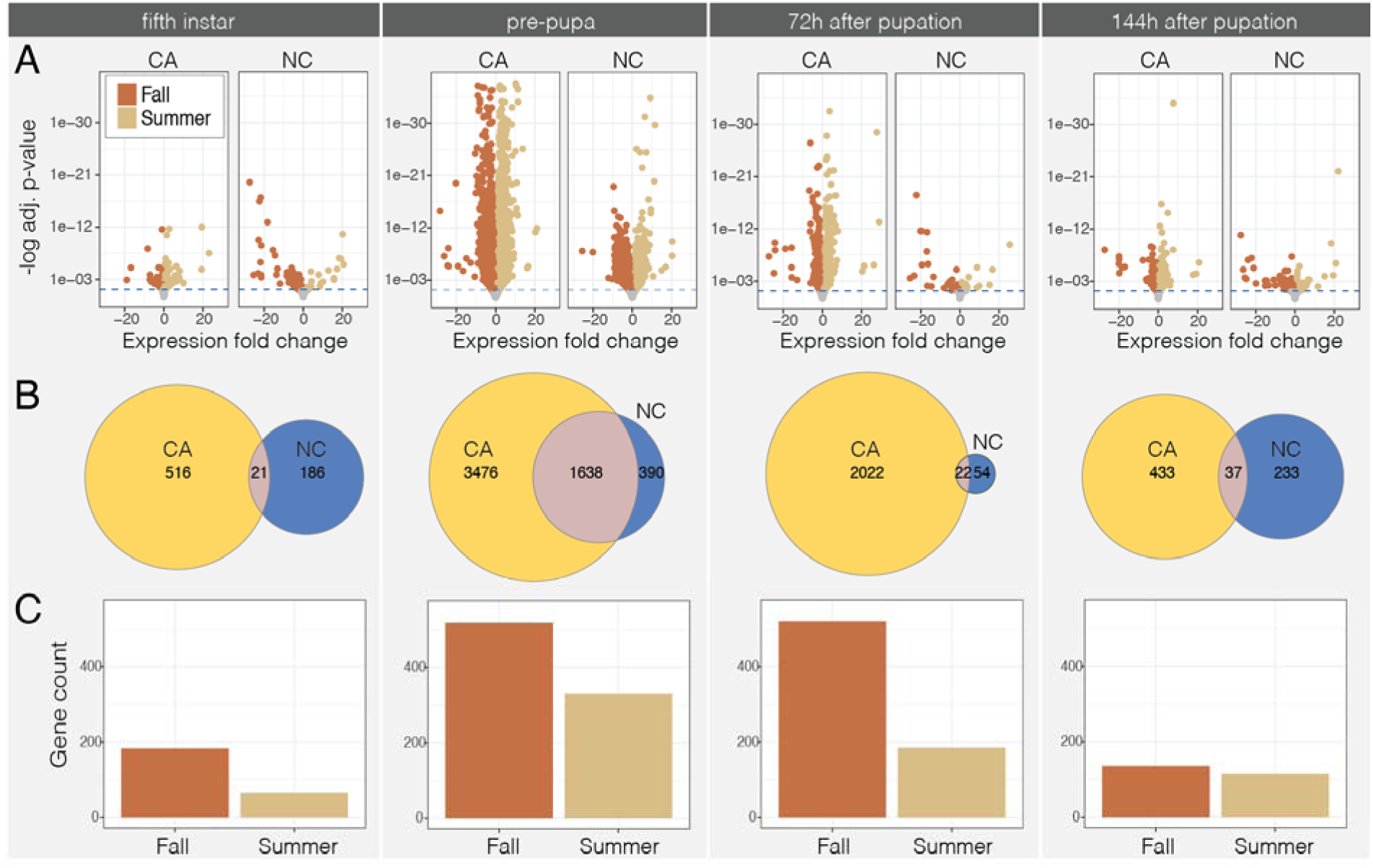
Gene expression plasticity is higher in a canalized population. A) Volcano plots of differentially expressed genes between rearing conditions (fall = red, summer = tan) across development in both California and North Carolina populations. B) Euler diagrams of the number of differentially expressed genes in each population and their overlap. C) The number of condition responsive genes at each stage of development (in either population) that are also differentially expressed between populations within each condition (fall = red, summer = tan). **Alt text**: Comparison of gene expression plasticity between California and North Carolina populations of *J. coenia*. A are volcano plots of differentially expressed genes between environmental treatments in each population and across four developmental stages. B are euler diagrams showing the overlap in the number of plastic genes between populations at each developmental stage sampled. C are bar plots showing the number of plastic genes that are also differentially expressed between populations, with one bar for each treatment.

**Figure 3.**
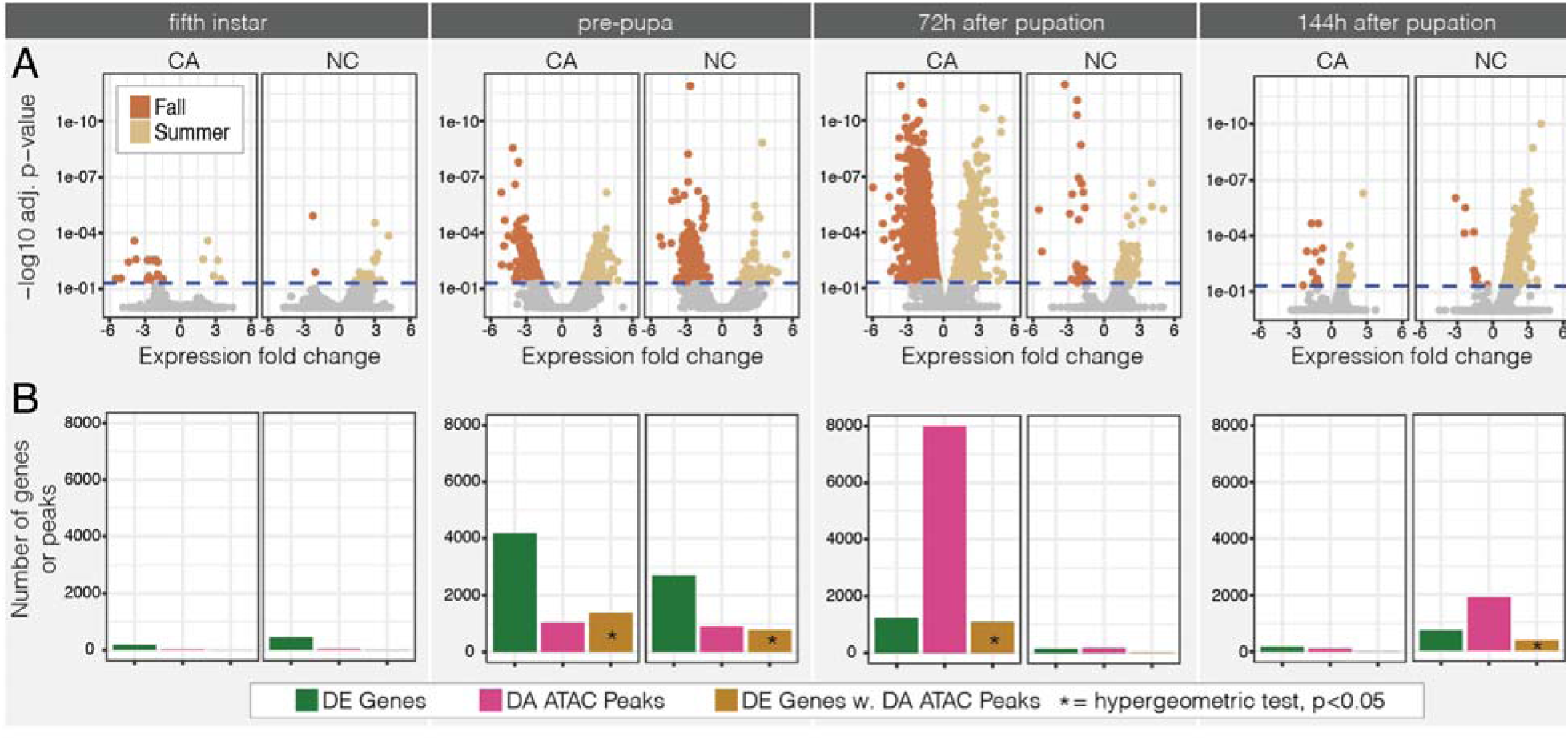
Chromatin accessibility is plastic and diverged between populations. A) Volcano plots of differentially accessible ATAC peaks between rearing conditions (fall = red, summer = tan) across development in both California and North Carolina. There is far more epigenetic plasticity during early pupal development in the California population than in North Carolina, but more epigenetic plasticity in late pupal development in North Carolina. The number of differentially expressed genes (green), differentially accessible ATAC peaks (pink), and the number of differentially expressed genes within 100kb of a differentially accessible ATAC peak (brown) across development. There are more DE genes with a differentially accessible (DA) ATAC peak than expected by chance in the pre-pupal stage of both populations, the early pupal stage in California and the late pupal stage in North Carolina. **Alt text**: Comparison of chromatin accessibility plasticity between populations and across four stages of development. A is a series of volcano plots of ATAC peak accessibility. B is a series of bar plots comparing the number of DE genes, differentially accessible (DA) ATAC peak, and the number of genes with a DA ATAC peak.

**Figure 4.**
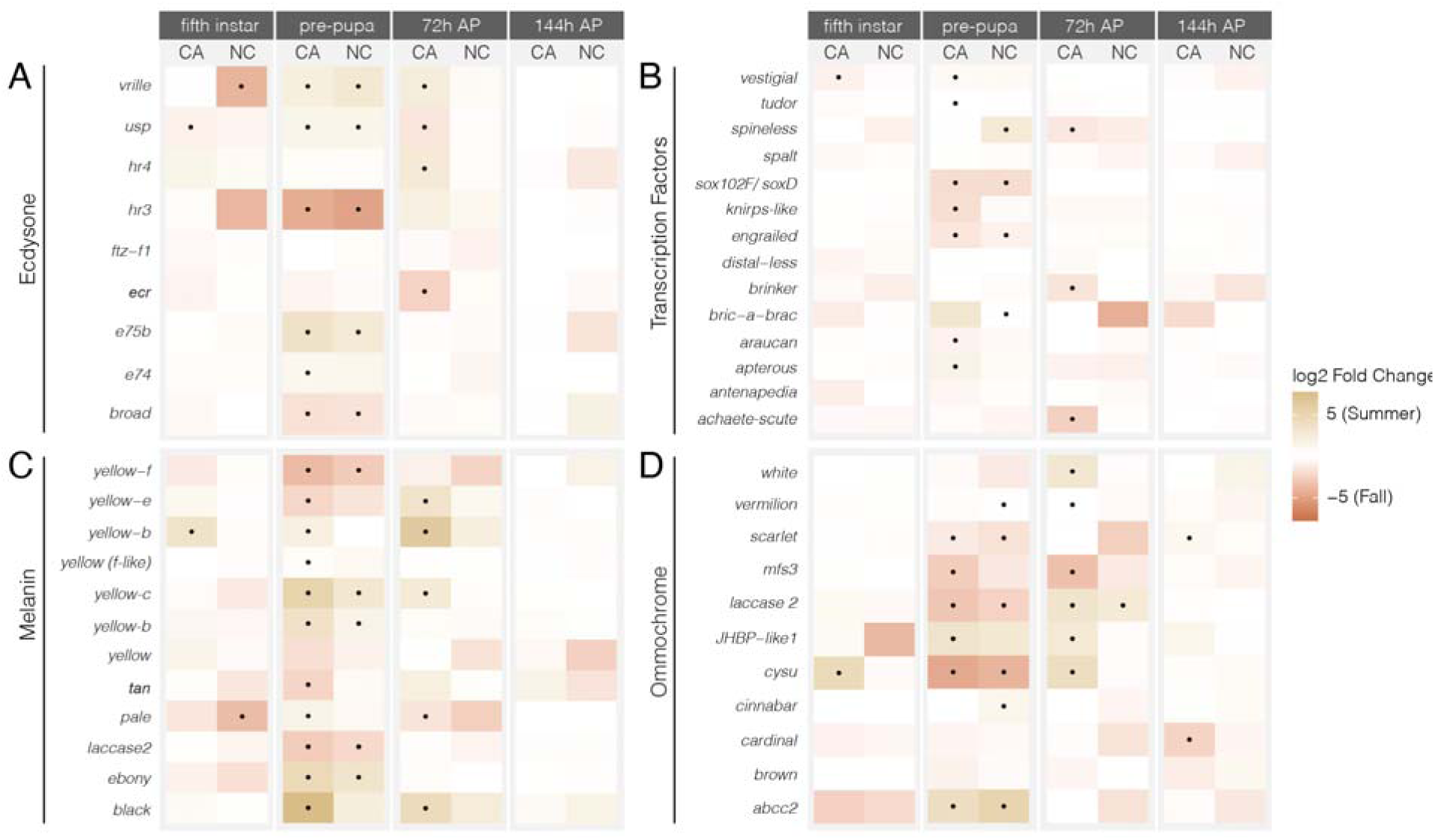
Key color pattern and endocrine signaling genes are plastic in the canalized population. Heatmaps of differentially expressed genes. A) ecdysone pathway, B) patterning genes C) melanin pathway, D) ommochrome pathway. Tan colors represent upregulation in summer-like conditions and red colors represent upregulation in fall-like conditions. Black dots indicate significantly differentially expressed genes. **Alt text**: Heatmaps comparing log2fold change of genes in four different categories: A: the ecdysone pathway, B: patterning transcription factors, C: the melanin pathway, and D: the ommochrome pathway.

To determine to what extent diverged transcriptional plasticity was caused by differences in how animals specifically respond to fall-like versus summer-like conditions, we compared gene expression between the two populations while controlling for environmental condition. We found that the difference in gene expression plasticity between North Carolina and California is driven more by changes to how animals respond to fall-like environmental conditions than by how animals respond to summer-like conditions (Fig. 2C).

### Epigenetic plasticity is greater, and heterochronically shifted, in a canalized population

We next asked whether divergence in an environmentally responsive epigenetic state, chromatin accessibility, is associated with the degree of morphological plasticity. Concurrent with the gene expression experiments described above, we quantified chromatin accessibility in both populations under summer-like and fall-like conditions at four stages of development. All samples passed QC for alignment and FRiP (Table S2). Samples generally clustered by developmental stage, though North Carolina 72h cold-reared sampled clustered more closely to earlier stages of development, suggesting that the epigenetic profile of the wing disc at this stage is shifted earlier in development as compared to the California population (Fig. S2). We found that the number of environmentally responsive ATAC peaks was highly dependent on developmental stage (Fig. 3A, Table S5, Table S6). During the late fifth instar there was relatively little epigenetic response to environmental conditions in either population (28 ATAC peaks in California; 44 ATAC peaks in North Carolina), while just 24 h later, during the pre- pupal stage, many more loci showed a chromatin accessibility response (1024 ATAC peaks in California; 893 ATAC peaks in North Carolina). During early pupal development, the number of differentially accessible ATAC peaks was much higher in animals from California versus North Carolina (7998 ATAC peaks in California; 176 ATAC peaks in North Carolina), but during late pupal development, that pattern reversed, and more ATAC peaks were differentially accessible in North Carolina animals (115 ATAC peaks in California; 1900 ATAC peaks in North Carolina). Thus, two surprising observations came from these experiments. First, the overall magnitude of epigenetic responsiveness was significantly greater in the morphologically canalized California population. Second, there appeared to be a timing shift in strength of the epigenetic response, where the California population showed most of its response early in pupal development, while North Carolina showed most of its response late in pupal development.

Next, we hypothesized that the observed divergence in gene expression plasticity between North Carolina and California *J. coenia* might be caused by variation in CRE activity near environmentally responsive genes. To evaluate whether plastic gene expression varied in association with environmentally induced changes in CRE activity, we used our ATAC-seq dataset to characterize the chromatin accessibility profiles of all genes with environmentally sensitive expression (Table S5, S6). We found that at the pre-pupal and late-pupal stages in North Carolina animals, and at the pre-pupal and early-pupal stages in California animals, there were more environmentally responsive genes adjacent to environmentally responsive ATAC peaks than expected by chance (Fig. 3B; hypergeometric test, p<0.05). This is consistent with expectations that environmentally induced changes in the epigenetic state of CREs play a direct role in transcriptional plasticity of a sizable subset of environmentally responsive genes.

### Plasticity of color patterning and endocrine gene expression varies between populations

As described above, we observed strong global patterns of gene expression plasticity in both California and North Carolina populations. We next wanted to assess whether known or suspected wing patterning genes may show differences in gene expression between populations and treatments. We thus characterized divergent responses to environmental cues between populations by evaluating changes in the expression of genes potentially involved in ecdysone signaling, wing pattern development, melanin pigmentation, and ommochrome pigmentation.

We began by performing an unbiased GO-term enrichment analysis to evaluate whether the abundance of plastic gene expression in the California population could be explained by a stress response to a novel environment. However, we did not observe an enrichment of stress-response genes in our analysis, suggesting that this is not the main source of plastic gene expression in the California population (Fig. S3).

We found that expression of ecdysone signaling genes was highly dependent on developmental stage and condition. The two genes that form the functional ecdysone receptor complex (*ecdysone receptor* and *ultraspiracle*) were significantly upregulated under fall-like conditions during early pupal development in Californian *J. coenia*, but not in the North Carolina population (Fig. 4A). For *ecdysone receptor*, and consistent with the genome-wide pattern, this divergence could be largely explained by a change in expression under fall-like conditions. The ecdysone-associated nuclear hormone receptor *hr4* was upregulated under summer-like conditions during early pupal development in the Californian population but not in the North Carolina population, a pattern also explained by a change in expression under fall-like conditions. The genes *e75b* and *hr3* were differentially expressed between conditions at the pre- pupal stage, but in opposite directions: *e75b* was upregulated under summer-like conditions while *hr3* was upregulated under fall-like conditions. Further, expression plasticity of *hr3* was reduced in North Carolina *J. coenia*. These observed expression plasticity divergences of key ecdysone pathway genes suggest potential candidates for upstream signal transducers that could drive larger scale differences between populations.

We next looked at candidate color patterning genes, as several of these exhibited environmentally induced expression changes at the pre-pupal and early pupal stages (Fig. 4B). Only two genes were condition sensitive at the same stage in both populations: *soxD* and *engrailed*. The environmental sensitivity of other genes has diverged; and many are environmentally modulated in only one population. Notably, the genes *spineless*, *brinker*, and *achaete-scute* were all upregulated under fall-like conditions during early pupal development in the California population and not in North Carolina. In contrast, *spineless* was upregulated in summer-like conditions while *bric-a-brac* was upregulated in fall-like conditions in North Carolina and not California. Therefore, among the many genes with expression plasticity, there is evidence that the environmental regulation of some potential color patterning factors has also diverged between populations.

In terms of melanin-associated genes, we were surprised to observe that the California population showed many more melanin-associated genes with environmentally responsive expression (11/17) than did the North Carolina population (5/17) (Fig 4C). These genes were slightly more likely to be upregulated under summer-like conditions than fall-like conditions, especially during early pupal development in California. Four genes, however, were condition sensitive in both populations at the pre-pupal stage: *ebony*, *laccase2*, *yellow-b*, and *yellow-c*, suggesting a core set of melanin genes may have a conserved response program across populations.

Many genes in the ommochrome biosynthesis pathway were found to be environmentally responsive (Fig 4D). For example, five genes in this pathway are upregulated under fall-like conditions in the California population at the pre-pupal stage and five genes are upregulated under summer-like conditions at the early pupal stage. Most of the environmentally responsive ommochrome genes at the pre-pupal stage were differentially expressed in both populations (4/6 genes), but *vermilion* and *cinnabar* were both environmentally responsive in North Carolina only, perhaps suggesting that these core ommochrome enzymes play an important functional role in the morphologically plastic population.

Evaluating all these candidate genes together, we found that their expression recapitulates what we observed generally across the genome: more genes have environmentally sensitive expression in the canalized California population compared the plastic North Carolina population. We found no support for global gene expression plasticity being driven by a stress response. We did, however, find that transducers of ecdysone signaling, patterning transcription factors, and pigmentation genes all have diverged plastic expression between populations.

### Environmentally responsive CREs are enriched for stage-specific transcription factors

We next sought to identify DNA sequence motifs associated with environmentally responsive ATAC peaks, reasoning that some of these motifs may be binding sites for transcription factors involved in coordinating developmental plasticity. We implemented a deep learning model that predicts accessibility at a base-pair scale and aggregated scores from this model into motifs(Shrikumar et al. 2018; Pampari et al. 2025). This method uncovered 72 motifs across all stages, treatments, and populations. Many of the motifs we identified showed significant similarity to *Drosophila* binding motifs, suggesting functional conservation (Fig. S5).

We next scanned for enrichment of any of these motifs in plastic ATAC peaks as compared to all ATAC peaks (Fig 5A). Within each population, the developmental stage largely determined which motifs were enriched. At the pre-pupal stage, 22 motifs were shared between populations, but at the subsequent two stages, there were very few overlapping motifs (Fig 5B; 72h = 0 motif, 144h = 6 motifs). However, there was substantial overlap (65 motifs) between the stage 72h after pupation in California and the stage 144h after pupation in North Carolina.

**Fig 5.**
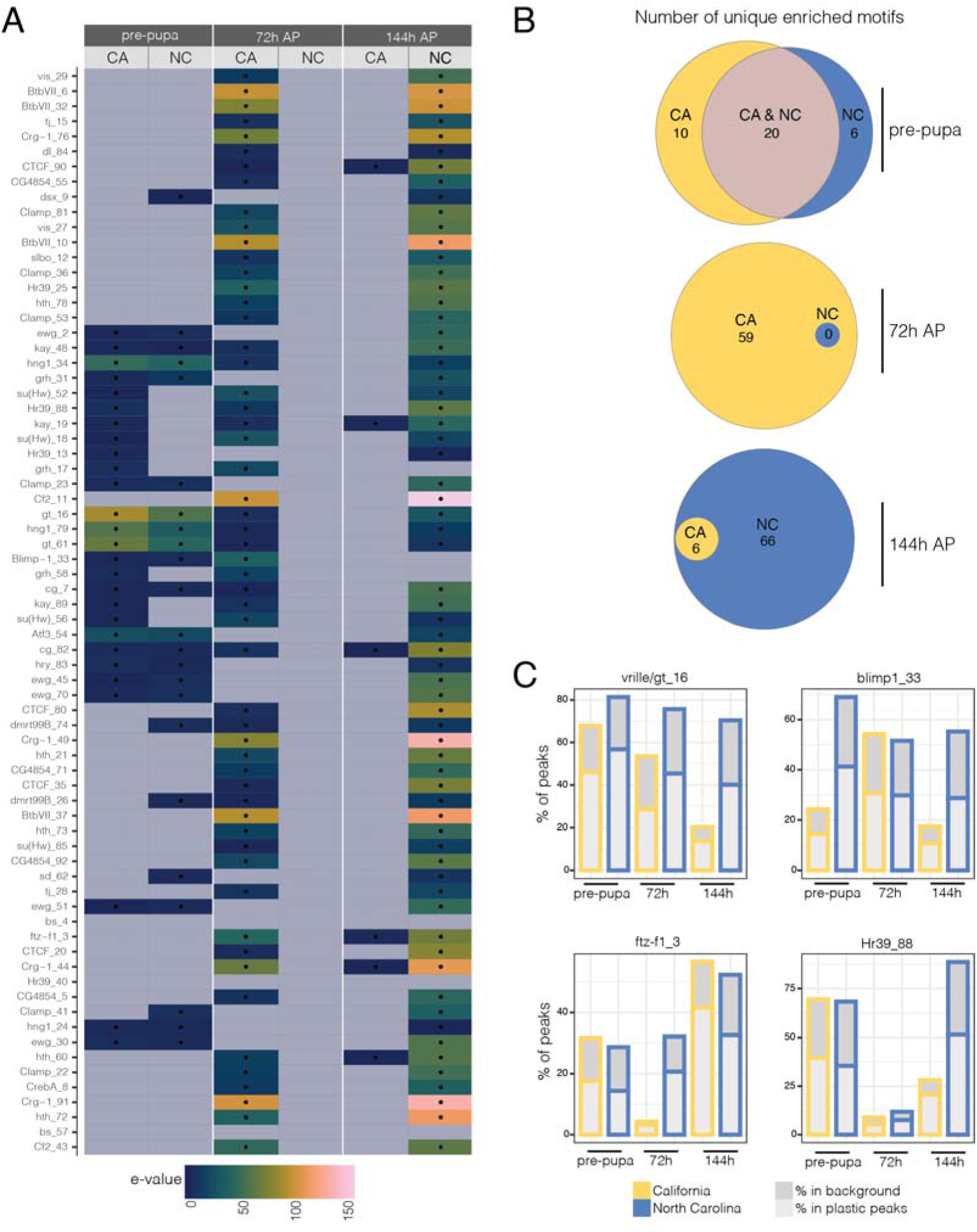
Motif enrichment analysis shows a heterochronic shift in plastic CREs. A) Heatmap of enrichment in plastic ATAC peaks compared to all ATAC peaks. B) Number of motifs that overlap between stages. C) Percentage of plastic ATAC peaks with specified motif across all stages and populations. **Alt text**: analysis of motifs enriched in plastic ATAC peaks. A is a heatmap of motif enrichment significance. B contains three euler diagrams showing the overlap between the number of enriched motifs between populations at each stage. C has four bar plots comparing the percentage of plastic ATAC peaks that contain a motif (for the most abundant motifs) between populations and across development.

The overlap of motifs enriched in plastic regulatory elements between an early stage in the California population (72h) and a later stage in the North Carolina population (144h) could be considered a heterochronic shift. Specifically, this suggests a group of transcription factors that regulate plastic chromatin accessibility has shifted activity to an earlier stage of development in the California population. *J. coenia* reared under fall-like conditions develop more slowly than their summer-like counterparts. Therefore, the plastic changes to chromatin accessibility that we observe could be caused by fall-like animals being sampled at a comparatively earlier stage of development than summer-like animals. However, both populations experience delayed development under fall-like conditions, so this likely does not explain the heterochronic shift we detect here.

Four of the most abundant motifs in plastic ATAC peaks correspond to transcription factors *ftz-f1*, *blimp1*, *hr39*, and *vrille* (this motif closely matched both *giant* and *vrille*) (Fig 5C). These genes are all known to be sensitive to ecdysone signaling and could therefore be important mediators of developmental plasticity. The expression of each of these genes is diverged between populations, and both *ftz-f1* and *vrille* are differentially expressed between environmental conditions as well (Fig. S5). We caution that these genes are examples of transcription factors that could bind the de-novo discovered motifs, as it is impossible to know the exact identity from these data. Lastly, we also found *ctcf* to be enriched in plastic ATAC peaks and to have diverged expression between populations under fall-like conditions. These motifs consistently associated with plastic ATAC peaks across the genome pinpoint candidate gene regulators for future consideration.

## Discussion

In this study we assessed how changes to gene expression and chromatin accessibility are associated with seasonal wing color plasticity in butterflies. The focal species of this study, *J. coenia*, displays strong wing color plasticity in some populations and very little plasticity in others, but the mechanistic basis of this variation remains unknown. Our expectation was that gene expression and epigenetic plasticity would be elevated in the population with greater morphological plasticity, and that we would identify environmentally responsive expression of genes involved in color patterning. To the contrary, we were surprised to find that the morphologically canalized population showed significantly greater transcriptional and epigenetic response to environmental conditions, and that numerous wing patterning candidate genes showed plastic response in both populations (Fig. 4A-D). We believe this counterintuitive case study may provide some insight into the evolution of plasticity mechanisms.

### How do we explain cryptic plasticity in gene regulation?

We observed extensive divergence in gene expression plasticity between populations of *J. coenia*, with the unexpected finding of significantly more environmentally sensitive gene expression throughout development in a population with a canalized, largely invariable phenotype. In other taxa, the opposite trend – a positive relationship between the extent of gene expression and phenotypic plasticity – has been observed (Alaux et al. 2009; Morris et al. 2014; Shaw et al. 2014; Casasa et al. 2020; Foquet et al. 2021). What might explain the unexpected pattern we observed? Broadly, we consider two alternative models: (1) *Developmental homeostasis*: the gene expression plasticity we observe in the canalized population plays a functional role in maintaining robust trait development, and (2) *Developmental masking*: the gene expression plasticity is largely non-functional and masked by one or more mutations that fix the trait, rendering it insensitive to environmental effects.

Homeostasis is an active process that requires the perception of an environmental condition and transduction to downstream targets that maintain a steady internal state. Genes that encode for heat shock proteins, for example, are upregulated during periods of environmental stress and stabilize protein conformation within cells (King and MacRae 2015). Likewise, plastic gene expression in California *J. coenia* could be a component of a mechanism that buffers color pattern development from environmental stimuli. If this is the case, we would expect plastic gene expression to occur earlier in development, either before or during the critical period when color pattern plasticity is induced. We also expect that the genes that respond to environmental cues in the canalized population would function to prevent those cues from being transduced in wing cells, such as by changing the expression of genes in the ecdysone response pathway.

In a developmental masking model, environmentally induced changes in gene expression have little or no effect on trait phenotype because they are masked by mutations that constitutively turn on the summer morph. Indeed, multiple studies have linked a small number of mutations downstream of ecdysone signaling to the evolution of a fixed fall morph in *J. coenia* (Rountree and Nijhout 1995; van der Burg et al. 2020). These mutations do not change the underlying processes that sense and signal environmental cues, but instead they buffer the final adult phenotype against responding to these cues. One consequence of developmental masking could be the accumulation of cryptic genetic variation, because when functional variation of a developmental system is not realized morphologically, mutations that arise in that system are sheltered from selection (Rutherford and Lindquist 1998; Rohner et al. 2013; Nijhout 2025). Masking, then, could have significant consequences for the evolution of plasticity: mutations that would otherwise be selected against could accumulate and then be exposed when the system is unmasked, acting like a capacitor that could generate novel phenotypes (Tawfeeq et al. 2024; Sayed et al. 2025).

### Environmental cue transduction has diverged between canalized and morphologically plastic populations

To evaluate our two proposed models, we sought to better understand the developmental mechanisms that underlie plastic morphological development and how they have diverged between populations of *J. coenia*. It is well documented that chromatin accessibility is dynamic throughout the development of many traits, driving spatial and temporal patterns of gene expression (Lyko et al. 2010; Niederhuber and McKay 2021; Lowe et al. 2022; Li et al. 2025). Importantly for this study, this is a key mechanism for transducing ecdysone cues to target tissues during *Drosophila* metamorphosis (Uyehara et al. 2017; Uyehara et al. 2022). Chromatin accessibility is also sensitive to environmental cues, and evolution has harnessed this flexibility as a mechanism for regulating the development of plastic traits (Ernst et al. 2015; Ecker et al. 2018; Lowe et al. 2022; Davidson and Moczek 2024)

Using ATAC-seq data, we found that chromatin accessibility is affected by environmental cues during *J. coenia* wing development. Furthermore, much of this epigenetic response has diverged between populations (Fig. 3A, S2). The developmental stage with the most divergence in plastic regulatory elements is early pupal development, when the concentration of ecdysone is highest in *J. coenia* reared under summer-like conditions. We also found that genes in the ecdysone pathway have diverged expression between populations: the *ecdysone receptor*, *ultraspiracle*, *hr4,* and *vrille* are differentially expressed between conditions in California during early pupal development. Because ecdysone signals are relayed, in part, by pioneer factors that modulate CRE accessibility, our data suggest that the transduction of environmental cues has diverged between populations of *J. coenia*, and the genes with diverged environmentally sensitive expression are likely factors that control this change.

There is also evidence that plasticity of regulatory element accessibility could cause changes in gene expression. By annotating regulatory elements with their neighboring genes, we found that most of the environmentally sensitive genes during early pupal development in California and late pupal development in North Carolina are near a CRE that changes accessibility in response to environmental cues (Fig. 3B). Therefore, the changes we found in the ecdysone transduction pathway could also be linked to our observation of greater environmentally sensitive gene expression in the canalized California population. Taken together, these data could lend support to our homeostasis model, because the diverged cue transduction mechanism could function to actively buffer the effect of the environment on color pattern development. However, quantitative changes in expression between the two populations do not prove that the function of the cue sensing and transduction mechanism has changed. Instead, it is possible that this mechanism is intact but blocked from effecting color pattern development.

Furthermore, while a stress response could be an important part of a homeostasis mechanism, we did not find evidence of this in the California population (Fig. S3), which led us to further consider the masking hypothesis.

### A temporal shift underlies divergence in epigenetic plasticity between populations

The divergence that we observed between populations in the epigenetic response to environmental treatments could be caused by differences in the expression of chromatin-remodeling genes. To identify the proteins that may regulate chromatin accessibility plasticity, we used a deep learning model (chromBPNet) to uncover sequence motifs enriched within environmentally responsive ATAC-seq loci (Shrikumar et al. 2018; Pampari et al. 2025). We were then able to identify candidate transcription factor binding sites within these sequence motifs, thus providing us with a list of candidate trans-factors that may modulate environmental responsiveness of chromatin.

While most of the enriched motifs at the pre-pupal stage were shared between the two populations, very few were shared between populations at the latter two stages (Fig 5A). Instead, we observed that most of the motifs enriched in the California population at the early pupal stage were activated 72hrs later in the North Carolina population. This heterochronic shift in transcription factor motif enrichment can partly be explained by the global shift in timing of chromatin accessibility we observed (Fig. 3B). Most interesting however, is that the majority of these motifs are the same between the two populations – the only difference was the developmental stage when they were environmentally sensitive. There are hundreds of transcription factors in the *J. coenia* genome, however, we recovered only a handful here that are enriched in environmentally responsive ATAC peaks. Therefore, our data suggest that the trans-factors associated with chromatin plasticity in *J. coenia* are largely the same between populations – the difference is that there is an overall timing shift in the epigenetic state of their targets between populations. Thus, we speculate that a heterochronic shift in epigenetic regulation is likely a major explanatory factor in the different plastic color pattern morphologies of the two populations.

Developmental heterochrony has been linked to color pattern variation in butterflies previously (Koch et al. 2000). Both a melanic mutant of *Papillio glaucus* and an eyespot mutant of *B. anynana* were linked to changes in the rate of wing scale development. In both species, colored pigments are deposited in scales earlier in development than melanic pigments (in eyespots, lighter colored scales emerge first), so when mutations push the rate of scale development in particular regions of the wing to be slower (*P. glaucus*) or faster (*B. anynana*), the pigments that emerge in these regions are darker or produce more eyespots, respectively. A similar mechanism could explain how the shift in timing of plastic changes to regulatory element accessibility alters color pattern development in *J. coenia*. In the North Carolina population, the set of transcription factors that are enriched in plastic ATAC peaks later in development could activate the switch to red pigments and large eyespots. In contrast, in the California population, the activity of these elements could be shifted outside of the window in which these phenotypes are determined, resulting in the canalized phenotype. This lends support to our masking hypothesis: by changing the timing of an environmental cue transduction mechanism, evolution stops the readout of this process (i.e. color development) sheltering the mechanism from selection. Furthermore, this places the focus on understanding *how* epigenetic chromatin remodeling is controlled, and more specifically, on how the timing of this process is regulated.

It is possible that pioneer factors regulate plastic chromatin accessibility in *J.* coenia and that variation between populations in the regulation of this factor controls a shift in its activity. We identified four transcription factors that could serve this function. The genes *ftz-f1*, *hr39*, *blimp1*, and *vrille* all have diverged expression between North Carolina and California and two of them (*vrille* and *hr39*) are also differentially expressed between environmental conditions (supplementary Fig. S5). These candidates stand out because they are all known to respond to ecdysone signals and because they are some of the most abundant motifs in plastic ATAC peaks (Fig 2.5C). Because these genes likely have conserved regulatory regions that connect them to ecdysone signaling, one or more of them could also regulate condition-sensitive chromatin remodeling. If this is the case, the heterochronic shift could be caused by changes to how these genes are regulated.

## Conclusion

The mechanisms that imbue traits with predictable responses to environmental cues are complex and involve interactions between multiple scales of biological organization. In this study, we explore these connections by looking at the relationship between epigenetic chromatin remodeling, gene expression, and wing color plasticity in *J. coenia* and how these processes have evolved between populations. Gene expression plasticity, surprisingly, was greater in a population with a canalized phenotype. Chromatin accessibility is very sensitive to environmental conditions, and we found a heterochronic shift in the identity of plastic CREs. We propose that these two processes are connected: that the heterochronic shift masks the gene expression plasticity by breaking the connection between environmental signals and downstream color development. This model could explain how plastic traits are rapidly tailored by evolution to meet local fitness demands.

### Data Accessibility and Benefit-Sharing

#### Data Accessibility

Raw sequencing data and associated metadata are available on NCBI GEO, via accessions: GSE342405 (RNAseq, access token: gpabmskifnitvet), GSE342850 (ATACseq, access token: urcdwagyrjqprgx). The code used to analyze these data are archived on Zenodo DOI: https://doi.org/10.5281/zenodo.21844506.

#### Benefit-Sharing

Not applicable

## Author contributions

K.R.L.v.d.B, R.D.R, and N.K.B conceived of the experiments. K.R.L.v.d.B. collected and reared animals and performed sequencing library preparation. N.K.B. analyzed the data and drafted the manuscript. R.D.R supervised the experiments and edited the manuscript.

## Acknowledgments

We thank Martik Chatterjee, Jeannie McDonald, Leslie Babonis, and Michael Sheehan for their helpful conversations and feedback related to this project. We thank the Cornell Biotechnology Resource Center for their guidance on sequencing and analysis methods. We also thank our dedicated team of undergraduate support staff for their help caring for the animals used in this study.

## Conflict of interest

The authors have no competing interests with this research.

## Ethics Statement

This study did not require ethics approval.

## Funding

This work was supported by National Science Foundation Grants IOS-2128164, IOS- 1753559, and IOS-1557443 to R.D.R as well as National Institutes of Health Grant 1R35GM161748-01 to R.D.R.. Any opinion, findings, and conclusions or recommendations expressed in this material are those of the authors(s) and do not necessarily reflect the views of the National Science Foundation.

## Notes

### Competing Interest Statement

The authors have declared no competing interest.

