## Supplemental Figures for "Canalization masks high levels of cryptic epigenetic and transcriptional plasticity in butterflies"

MOLECULAR ECOLOGY

### Table of Contents:

| | |
|---|---|
| **Figure S1** | Page 2 |
| **Figure S2** | Page 3 |
| **Figure S3** | Page 4-7 |
| **Figure S4** | Page 8-13 |
| **Figure S5** | Page 14 |

MOLECULAR ECOLOGY

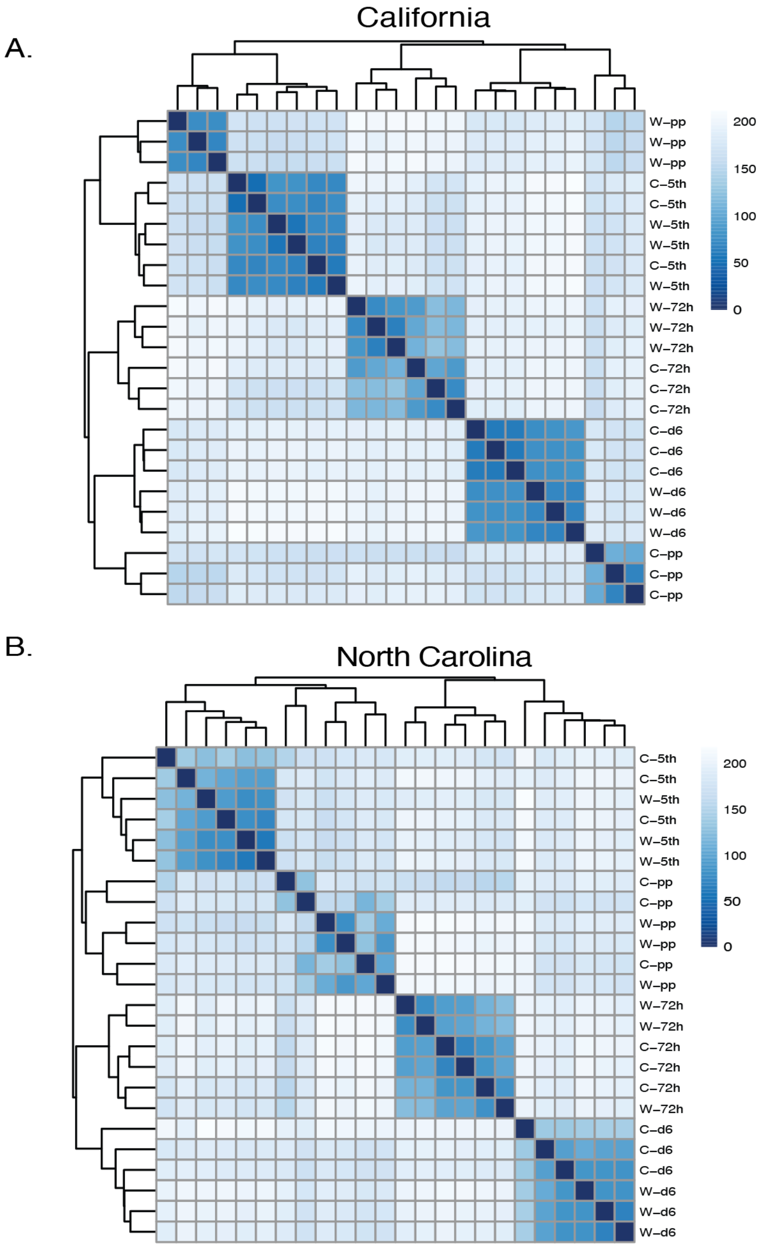


**Supplementary Figure S1.** RNA-seq sample distance heatmap. Samples generally cluster by stage and temperature treatment: w = warm, c = cold, $5^{th}$ = $5^{th}$ instar, pp = pre-pupal, 72h = 72h post pupation, d6 = 144h post pupation.

MOLECULAR ECOLOGY

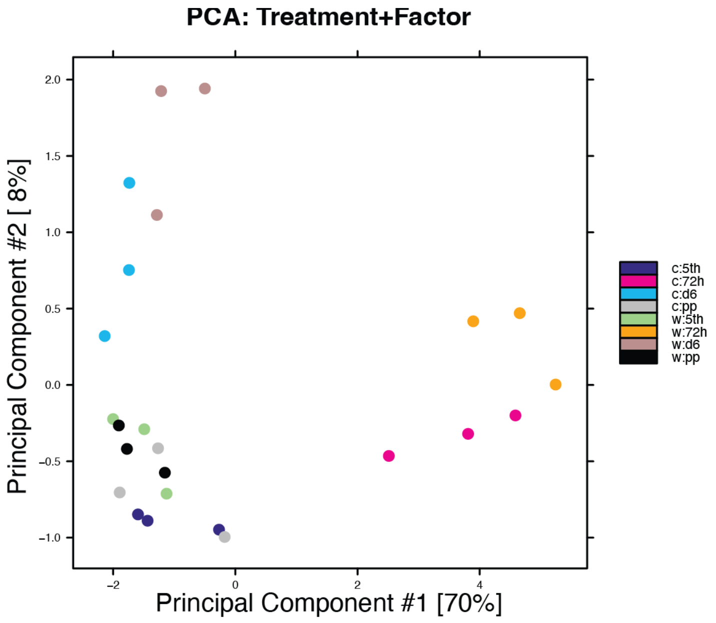


B.

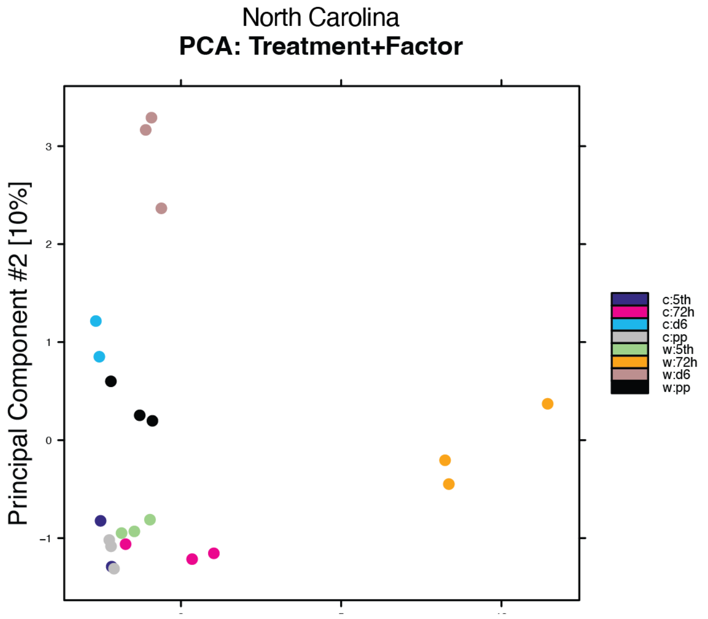


**Supplementary Figure S2.** ATAC-seq sample PCA. Samples generally cluster by stage and temperature treatment, except for North Carolina Cold 72h: w = warm, c = cold, 5th = 5th instar, pp = pre-pupal, 72h = 72h post pupation, d6 = 144h post pupation.

MOLECULAR ECOLOGY

Enriched GO terms across stages in CA

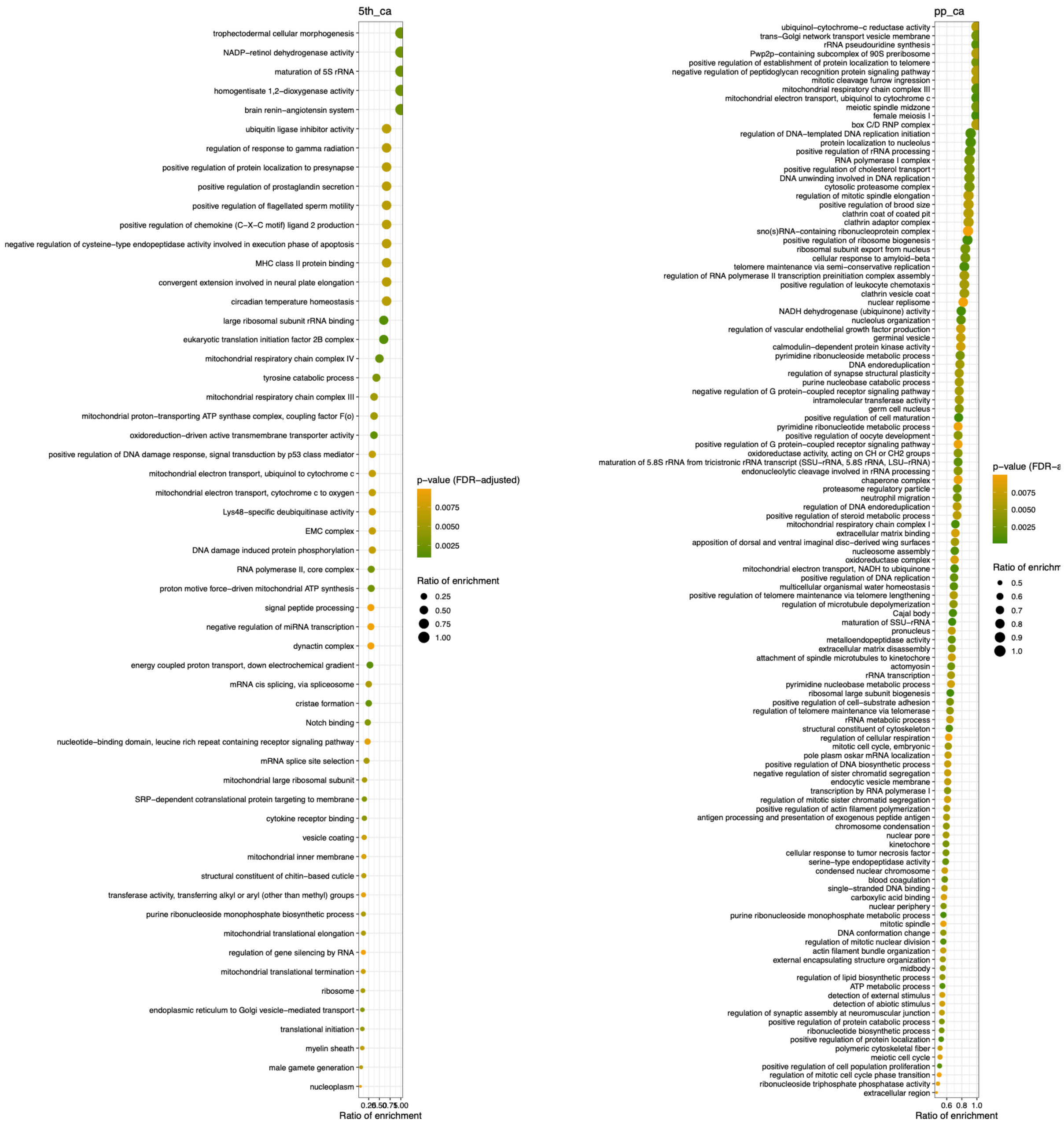

MOLECULAR ECOLOGY

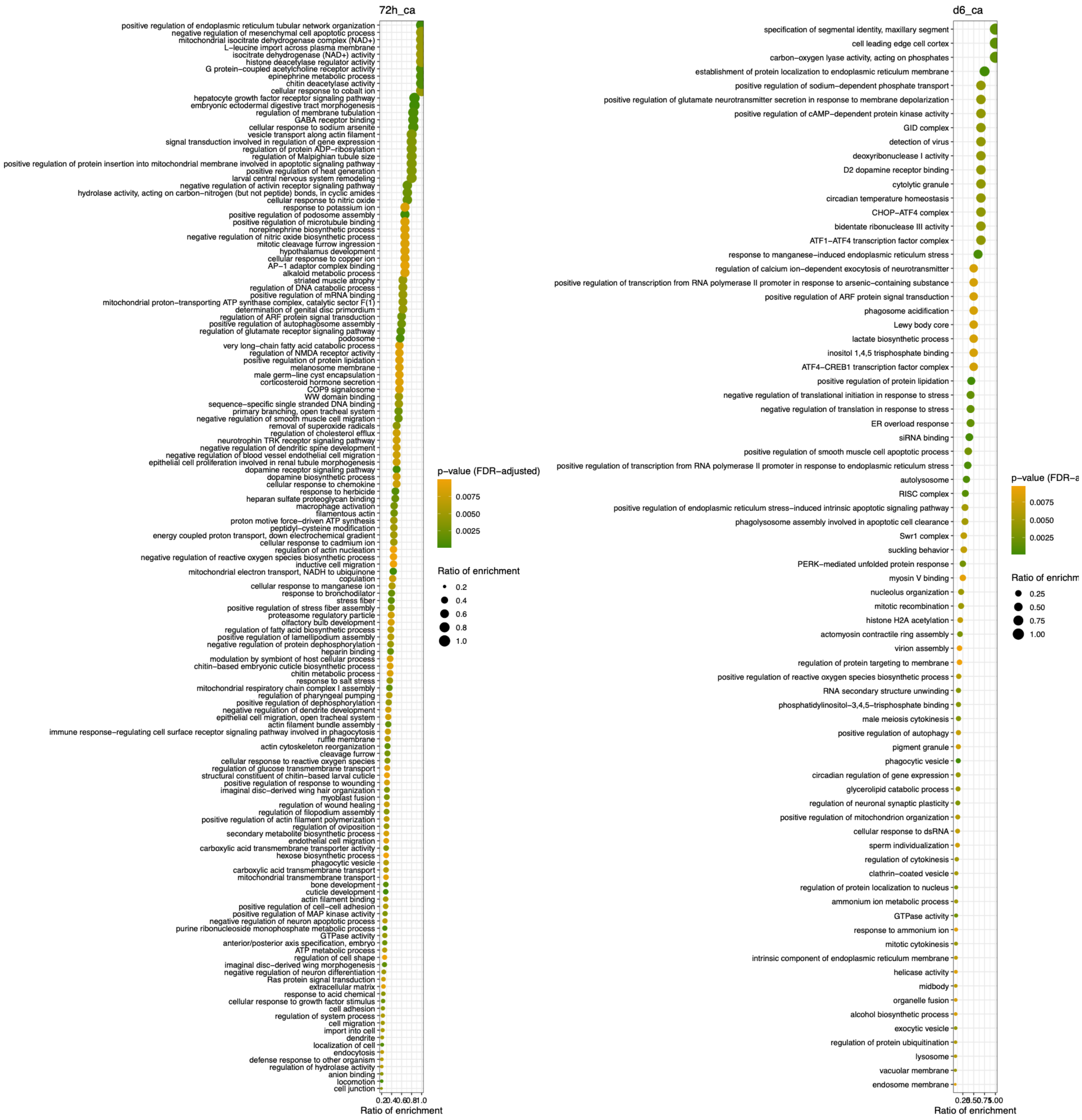

MOLECULAR ECOLOGY

Enriched GO terms across stages in NC

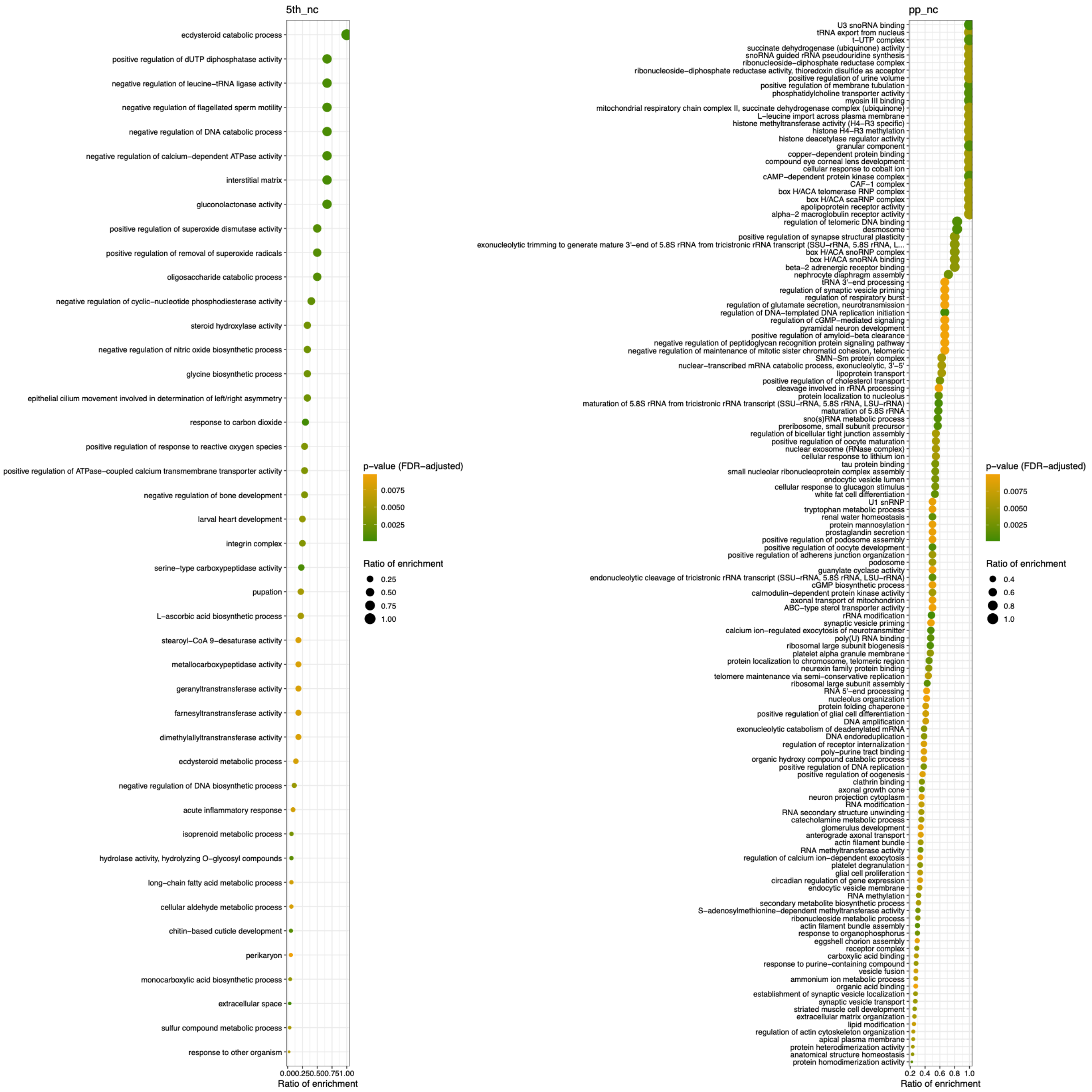

MOLECULAR ECOLOGY

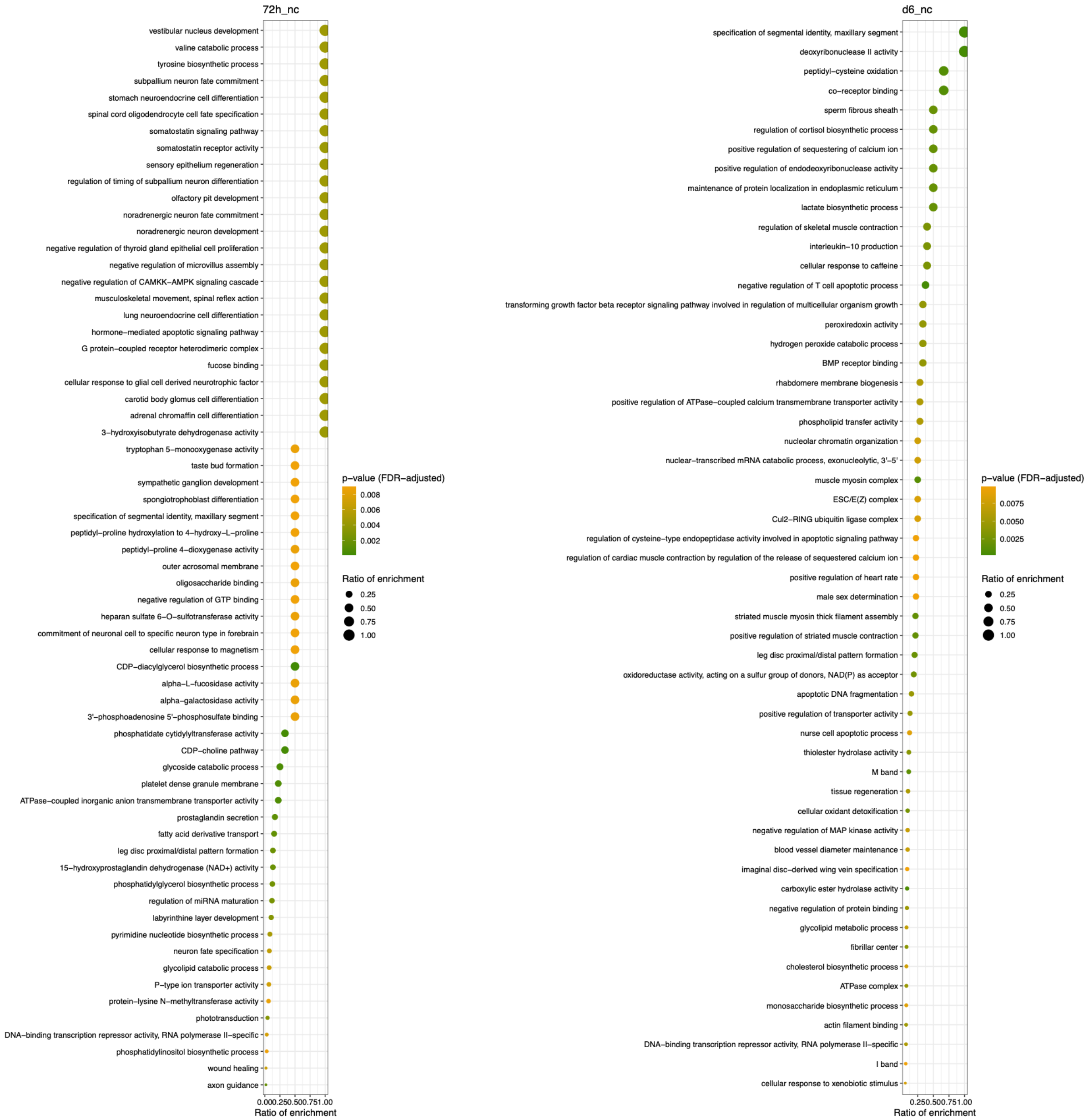


**Supplementary Figure S3.** Plots of GO term enrichment in differentially expressed genes at each stage of development in both California and North Carolina. $5^{th}$ = $5^{th}$ instar, pp = pre-pupal, 72h = 72h post pupation, d6 = 144h post pupation.

MOLECULAR ECOLOGY

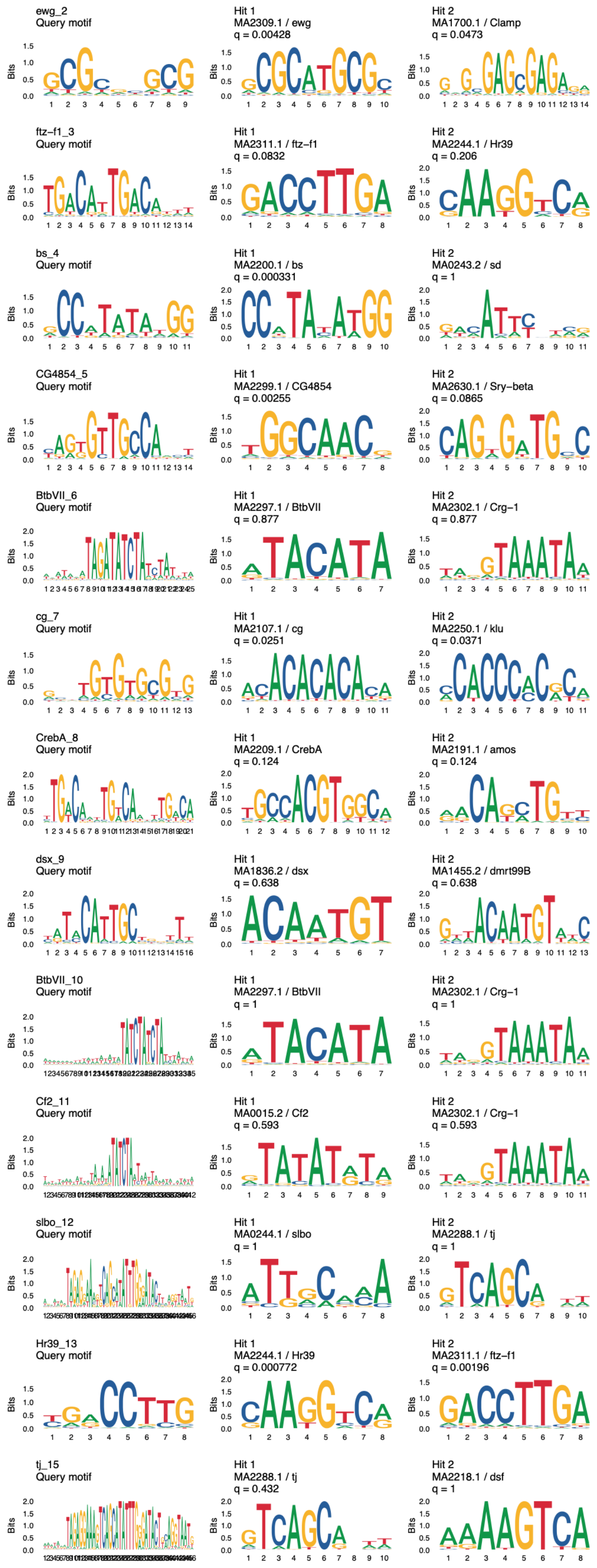

MOLECULAR ECOLOGY

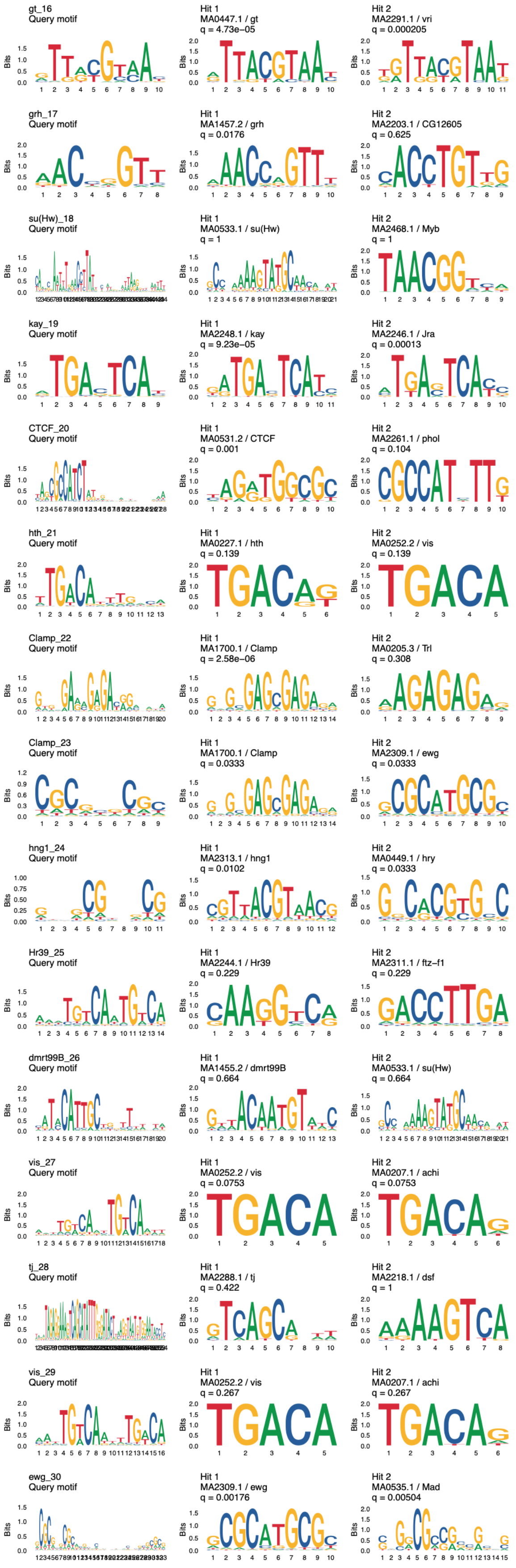

MOLECULAR ECOLOGY

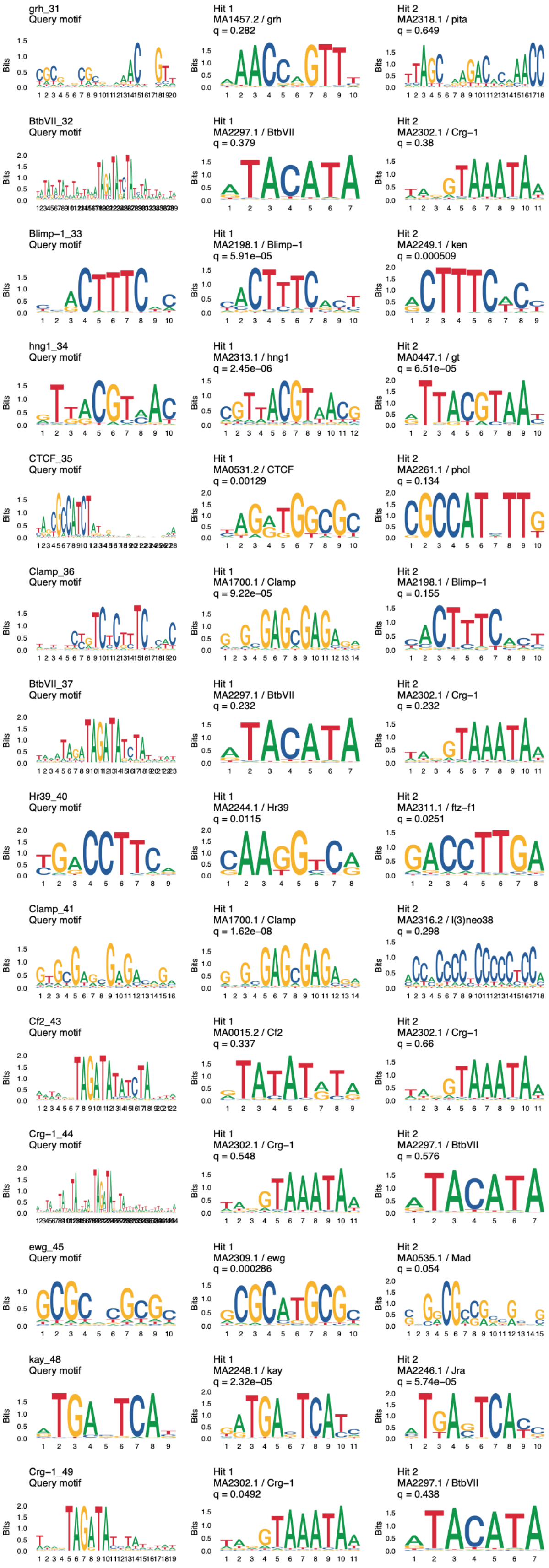

MOLECULAR ECOLOGY

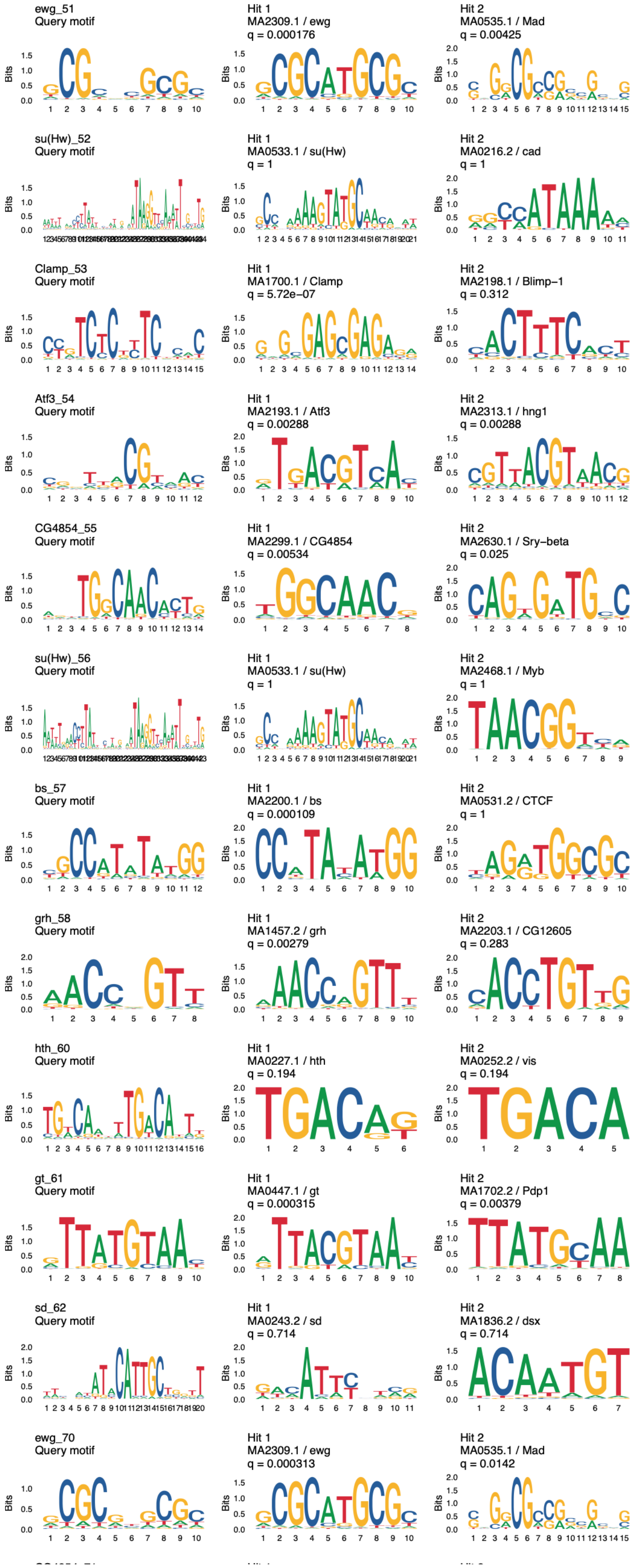

MOLECULAR ECOLOGY

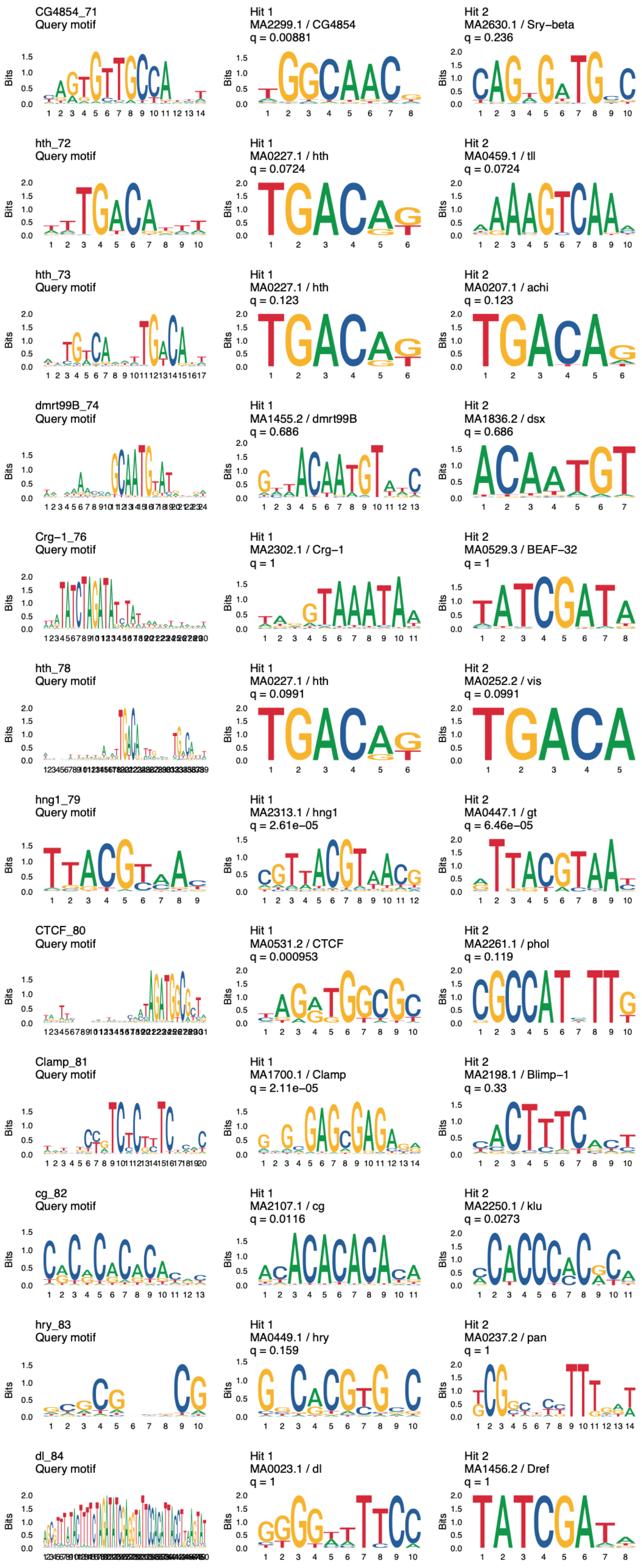

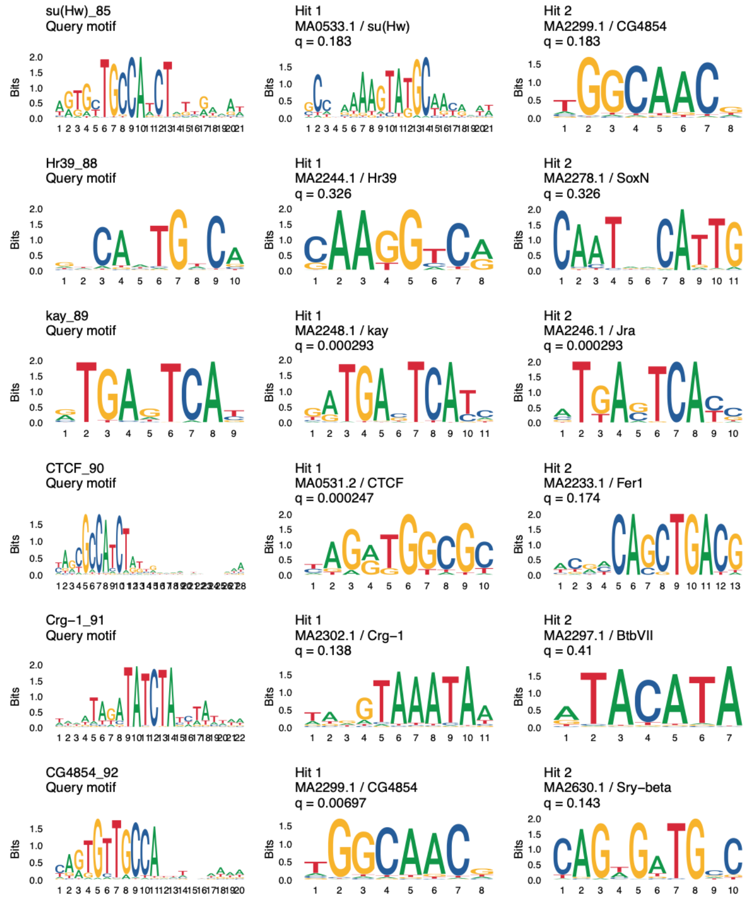


**Supplementary Figure S4.** De-novo discovered motifs (first column) across all samples and top two TomTom matchs (second two columns).

MOLECULAR ECOLOGY

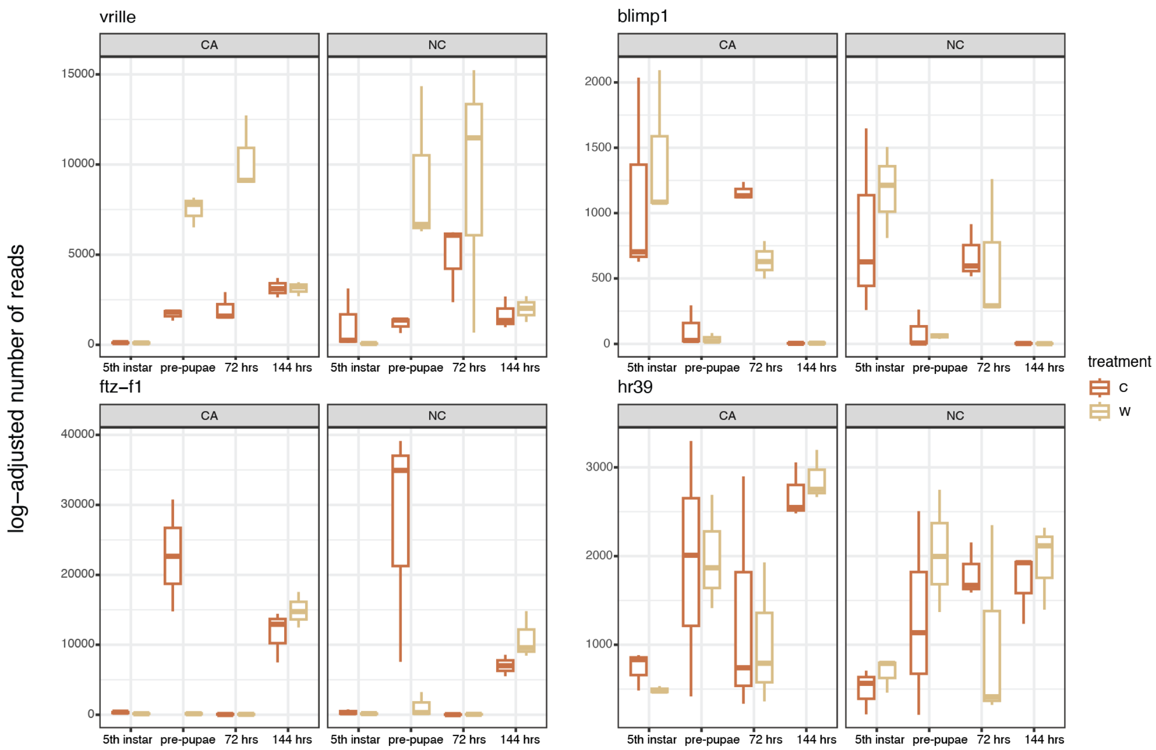


**Supplemental Figure S5.** Read count boxplots of transcription factors with the most abundant motifs in differentially accessible ATAC peaks.
